# Noncanonical PI(4,5)P2 coordinates lysosome positioning through cholesterol trafficking

**DOI:** 10.1101/2025.01.02.629779

**Authors:** Ryan M. Loughran, Gurpreet K. Arora, Jiachen Sun, Alicia Llorente, Sophia Crabtree, Kyanh Ly, Ren-Li Huynh, Wonhwa Cho, Brooke M. Emerling

**Author notes:** Corresponding author (B.M.E.).

## Abstract

In p53-deficient cancers, targeting cholesterol metabolism has emerged as a promising therapeutic approach, given that p53 loss dysregulates sterol regulatory element-binding protein 2 (SREBP-2) pathways, thereby enhancing cholesterol biosynthesis. While cholesterol synthesis inhibitors such as statins have shown initial success, their efficacy is often compromised by the development of acquired resistance. Consequently, new strategies are being explored to disrupt cholesterol homeostasis more comprehensively by inhibiting its synthesis and intracellular transport. In this study, we investigate a previously underexplored function of PI5P4Ks, which catalyzes the conversion of PI(5)P to PI(4,5)P_2_ at intracellular membranes. Our findings reveal that PI5P4Ks play a key role in facilitating lysosomal cholesterol transport, regulating lysosome positioning, and sustaining growth signaling via the mTOR pathway. While PI5P4Ks have previously been implicated in mTOR signaling and tumor proliferation in p53-deficient contexts, this work elucidates an upstream mechanism that unifies these earlier observations.

## Introduction

Phosphatidylinositol (PI) signaling is a diverse cellular process regulated by various phosphatases and kinases localized to specific cellular compartments. The phosphoinositide (PIP) family is comprised of seven phosphorylated derivatives of PI, which are localized to and embedded into specific intracellular membranes (*1, 2*). The PI composition of each membrane is highly regulated and site-specific, and unique PI signatures have been observed at different organellar membranes (*3–5*). In turn, the composition of phosphoinositides at various membrane compartments provides spatiotemporal cues directing membrane dynamics and determines protein recruitment to these sites (*6*). Although the most well-studied PIP modality is centric to the PI3K/AKT/mTOR signaling pathway (*7–9*), lesser-known PI-mediated signaling pathways are also under active investigation. Of recent interest is the role of PI5P4K in supporting tumor metabolism (*10–12*). In mammals, the phosphatidylinositol 5-phosphate 4-kinases (PI5P4Ks) are comprised of three isoforms (PI5P4Kα, PI5P4Kβ, PI5P4Kγ) responsible for the non-canonical generation of phosphatidylinositol 4,5 bisphosphate (PIP2) from phosphatidylinositol 5-monophosphate (PI(5)P).

To date, several relevant findings have shed light on the importance of PI5P4Ks maintaining cellular homeostasis under stress. Our original work showed that suppression of the most catalytically active PI5P4K isoforms (ɑ and β) in *TP53* deficient cancer cells inhibits proliferation, and the deletion of these enzymes in *Trp53* knockout mice confers protection from tumorigenesis (*13*). Further studies have revealed that PI5P4Ks localize to lysosomes and are necessary for autophagosome-lysosome fusion during stress in the p53-deficient context (*14*). Recently, the generation of PIP2 at the peroxisomal membrane by PI5P4Kɑ has been shown to facilitate the handoff of lysosomal cholesterol to the peroxisome, demonstrating a key role for these enzymes in the dynamics of organelle crosstalk (*15, 16*). Taken together, these findings suggest a critical role for PI5P4Ks in maintaining metabolic homeostasis by lysosome-related mechanisms.

Notably, the lysosome has emerged as an essential metabolic signaling organelle due to its several roles in nutrient sensing that lead to the translocation of mTORC1 to the lysosomal surface for activation (*17, 18*). When cells are supplied with sufficient nutrients, mTORC1 is recruited and anchored to the lysosomal surface by Rag GTPases. The Rag GTPases function as amino acid sensors in which sufficient levels of amino acids promote Rag heterodimer binding to mTORC1. Once anchored, mTORC1 encounters the Rheb GTPase, which triggers the kinase activity of mTORC1 and enables substrate phosphorylation. Intriguingly, novel mechanisms of cholesterol-dependent mTORC1 signaling are now being uncovered at the lysosome. Notably, these consist of lysosomal transmembrane proteins harboring cholesterol sensing domains, which have been shown to have independent mTORC1 activation mechanisms distinct from amino acid sensing (*19–22*). These findings present new questions about the importance of lysosomal cholesterol balancing, especially in the context of p53-deficient cancers, in which the negative regulation of feedback from high cholesterol can be circumvented via loss of p53 function. Indeed, cellular cholesterol content is tightly regulated by the abundance of cholesterol itself, where binding at sterol sensing domains retains SREBP-2 at the membrane of the ER, thus preventing transcriptional activation of the mevalonate pathway target genes (*23, 24*). Thus, proteins that prevent or promote lysosomal cholesterol efflux could have significant importance for future targeted therapies. In this study, we elucidate the molecular mechanisms underlying the role of the PI5P4Ks in lysosomal cholesterol transport and lysosome function in the context of p53-deficiency.

### PI5P4Ks are critical for tumor progression and rarely mutated in human cancer

Recent findings underscore the crucial role of PI5P4Ks in cancer progression and cellular homeostasis under stress (*13, 25*). We have previously shown that the catalytically active PI5P4K isoforms, ɑ and β, are vital for neonatal survival by enabling the completion of autophagy, a process impaired by their loss (*14*). In mice with a germline deletion of *Trp53*, two alleles of *Pip4k2a*, and one allele of *Pip4k2b*, tumor initiation was markedly suppressed, revealing a targetable vulnerability within the *Trp53* deficient background (*13*). However, deletion of all four alleles in adult animals has not yet been investigated for mouse survival and protection from tumorigenesis. To accomplish this, we generated a model for conditional and systemic deletion of p53 and both PI5P4K isoforms (*Pip4k2a*^flx/flx^ *Pip4k2b*^-/-^ *Trp53*^flx/flx^) crossed with a tamoxifen-inducible Cre-ERT2 fusion gene under the control of the human ubiquitin C promoter (*Ubc-cre+)* (*26*). At 8 to 12 weeks of age, *Pip4k2a^flx/flx^ Pip4k2b^-/-^ Trp53^flx/flx^ Ubc-cre+* mice were treated with tamoxifen to induce recombination in a temporal manner (Fig. 1A, Fig. S1A). As a control, p53 knockout mice with functional PI5P4Ks, *Trp53*^flx/flx^ *Ubc-cre+*, were treated simultaneously. Within 4-6 months of tamoxifen treatment, the p53 knockout mice with functional PI5P4Ks had developed tumors and required sacrifice (Fig. 1A). However, mice with full deletion of PI5P4Ks and loss of p53 exhibited complete protection from tumorigenesis up to 12 months post-treatment (Fig. 1A).

**Fig. 1.**
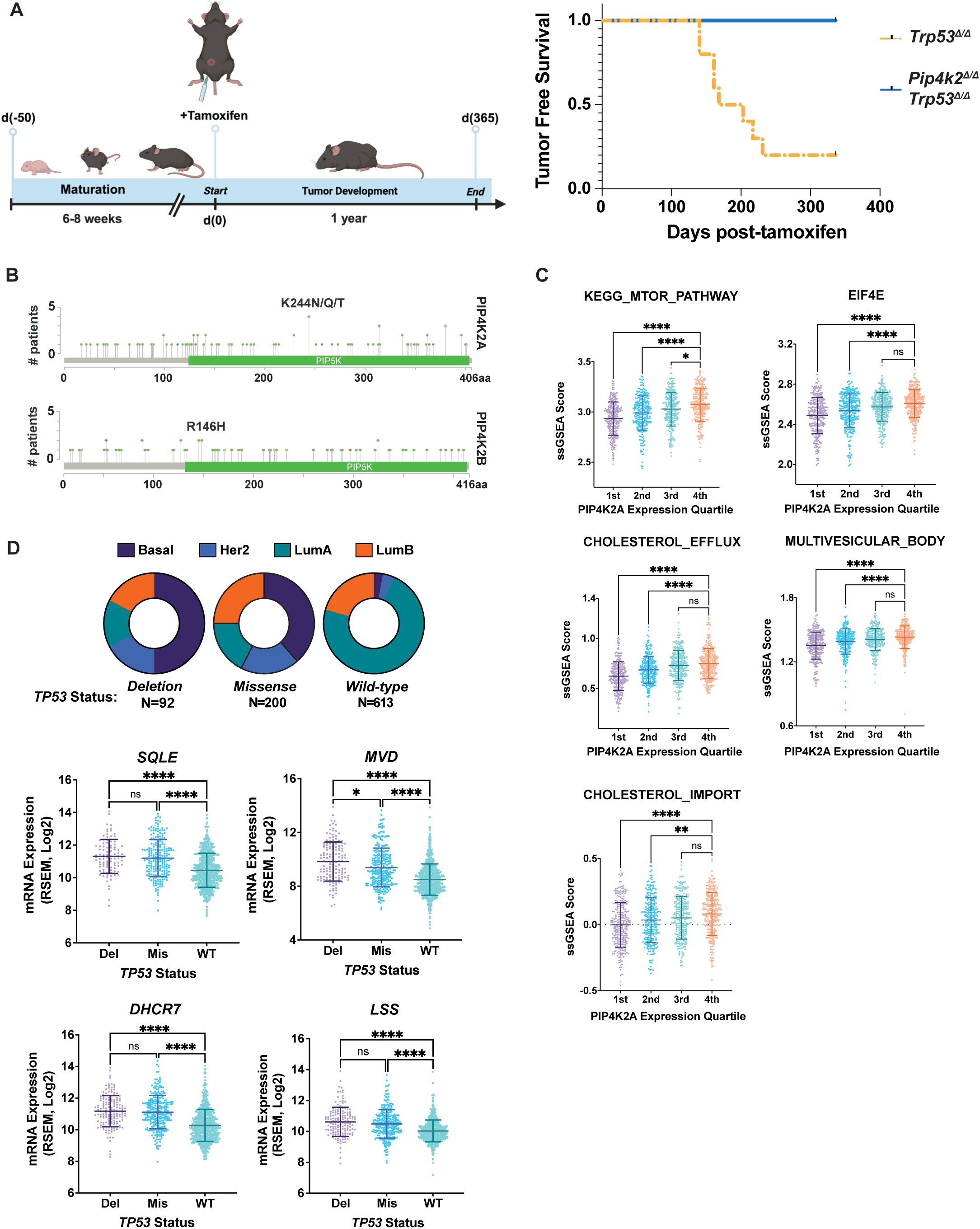
PI5P4Ks are crucial for support of tumor maintenance and are rarely mutated in human cancer. **(A)** *(left)* Schematic of tamoxi-fen-inducible recombination to generate knockout animals and monitor tumorigenesis (*Pip4k2aflx/flx Pip4k2b-/- Trp53flx/flx Ubc-cre+* vs *Trp53flx/flx Ubc-cre+*). *(right)* Tumor free survival of *Trp53-/-* vs. *Pip4k2a-/- Pip4k2b-/- Trp53-/-* mice post-tamoxifen mediated recombination. N=8 mice/group. Log-rank test, p=***0.0002, median tumor-free survival of *Trp53-/-* =185.5 days post-tamoxifen. **(B)** *PIP4K2A* and *PIP4K2B* mutations in TCGA pan-cancer analysis. Lollipop plot represents number of patients per mutation site. Total patient samples n=10,967. **(C)** GSVA analysis of TCGA BRCA dataset. Quartile RSEM values used for expression of *PIP4K2A*: 1st < 2nd < 3rd < 4th. N=270 patients per quartile. Significance was tested using Kruskal-Wallis test. p<**0.01, ***0.001, ****0.0001. **(D)** TCGA BRCA patients grouped by PAM50 subtypes. RNA-seq expression data of mevalonate pathway genes. Samples are segregated by *TP53* deletion, missense mutation, or wild-type in cBioPortal. Significance was tested using Kruskal-Wallis test. p<**0.01, ***0.001, ****0.0001..

The PI5P4Ks are highly conserved enzymes from metazoans to mammals. Transcript expression levels of *PIP4K2A*, *PIP4K2B*, and *PIP4K2C* have been profiled across all human cancers and show variable expression, suggesting that tissue context-specific metabolic profiles are likely to have a role in the requirement of these kinases (*11*). Mutation analysis of both PI5P4Kɑ and PI5P4Kβ across all cancers using The Cancer Genome Atlas (TCGA) pan-cancer datasets reveals that mutations of these enzymes are exceedingly rare, with little to no common mutational occurrences across all cancers (*27–29*) (Fig. 1B, Fig. S1B). The lack of mutations would suggest that p53-related chromosomal instability or amplification of PI5P4Ks is neither beneficial nor detrimental to the growth of p53-deficient cancers. Although rarely mutated, the PI5P4Ks have been shown to have elevated expression in breast cancer, using transcriptomics and immunohistochemistry of patient samples (Fig. S1D) (*12, 13*).

To uncover associations of the *PIP4K2A* gene expression with gene set enrichment (GSE) pathways, we utilized gene set variation analysis (GSVA) using the TCGA-BRCA dataset (*30*). Upon subdividing normalized *PIP4K2A* expression values into quartiles, we performed pathway analysis and found that increased gene expression correlated with autophagy, mTOR activation, and cholesterol-specific trafficking gene pathways (Fig. 1C). Previous work has implicated PI5P4Kα in cholesterol diffusion from the lysosome to the peroxisome (*15, 16*). However, the importance of these enzymes in cholesterol trafficking from the lysosome has yet to be explored in the p53-deficient context, where cholesterol biosynthesis is enhanced by coactivation of SREBP-2 and loss of p53 transactivational activity (*31–33*). Analysis of patient breast cancer samples by clinical subtype identifiers reveals that subtypes harboring a greater percentage of TP53 deletions or mutations often coincide with enhanced expression of the mevalonate pathway (Fig. 1D, Fig. S1C, Fig. S1E). Upon subdividing patient samples by *TP53* deletion, mutation, or wild-type status, we found enhanced transcriptional expression of several mevalonate biosynthesis genes (Fig. 1D, Fig. S1D). Together, these findings reinforce the importance of the PI5P4Ks for p53-deficient tumor maintenance and provide evidence that the PI5P4Ks are linked to cholesterol balance.

### PI5P4Ks support cholesterol homeostasis through lysosome positioning and mTOR activation

To explore the supportive role of the PI5P4Ks in cholesterol homeostasis when p53 function is compromised, we generated mouse embryonic fibroblasts (MEFs) from our conditional *Trp53* knockout models (*Pip4k2a-/- Pip4k2b-/- Trp53-/-* (p53-/- αβ^DKO^) and *Pip4k2a+/+ Pip4k2b-/- Trp53-/-* (p53-/- WT) control (Fig. S2A). To first measure cholesterol abundance in the PI5P4K-deficient context, we used the polyene antibiotic filipin as a marker for cellular free cholesterol. We observed a significant increase in free cholesterol in the p53-/- αβ^DKO^ MEFs compared to the control, which could be rescued upon re-expression of functional mouse PI5P4K⍺ (p53-/- α^WT^), but not with the expression of the catalytically-dead PI5P4K⍺ mutant (p53-/- α^KD^) (Fig. 2A). As a control for lysosomal cholesterol accumulation, cells were treated overnight with the amphiphilic cholesterol transporter U18666A, which disrupts the major cholesterol exporter Niemann-Pick Type C1 (NPC1) (*34*). This inhibition of cholesterol efflux from the lysosome had an additive effect of increased cholesterol in the p53-/- αβ^DKO^ and p53-/- PI5P4K⍺^KD^ when compared to p53-/- WT and p53-/- PI5P4K⍺^WT^ re-expression (Fig. 2A). To determine whether cholesterol accumulation occurred in the lysosomal compartment, we performed co-staining of filipin with lysotracker to mark the late endosome/lysosome compartment. In the PI5P4K-deficient context, we observed a significant increase in filipin overlapping with lysotracker, suggesting most cellular free cholesterol was contained within the lysosome (Fig. 2B). In addition, RNA-sequencing of mouse liver tissue from our previous work revealed that targeted deletion of PI5P4Ks in the hepatocytes significantly impaired cholesterol homeostasis and mTORC1 signaling (*35*) (Fig. S2B). These findings demonstrate the importance of PI5P4K function in the efflux of lysosomal cholesterol, a critical process in cells with enhanced cholesterol biosynthesis, such as p53-deficient tumors.

**Figure. 2.**
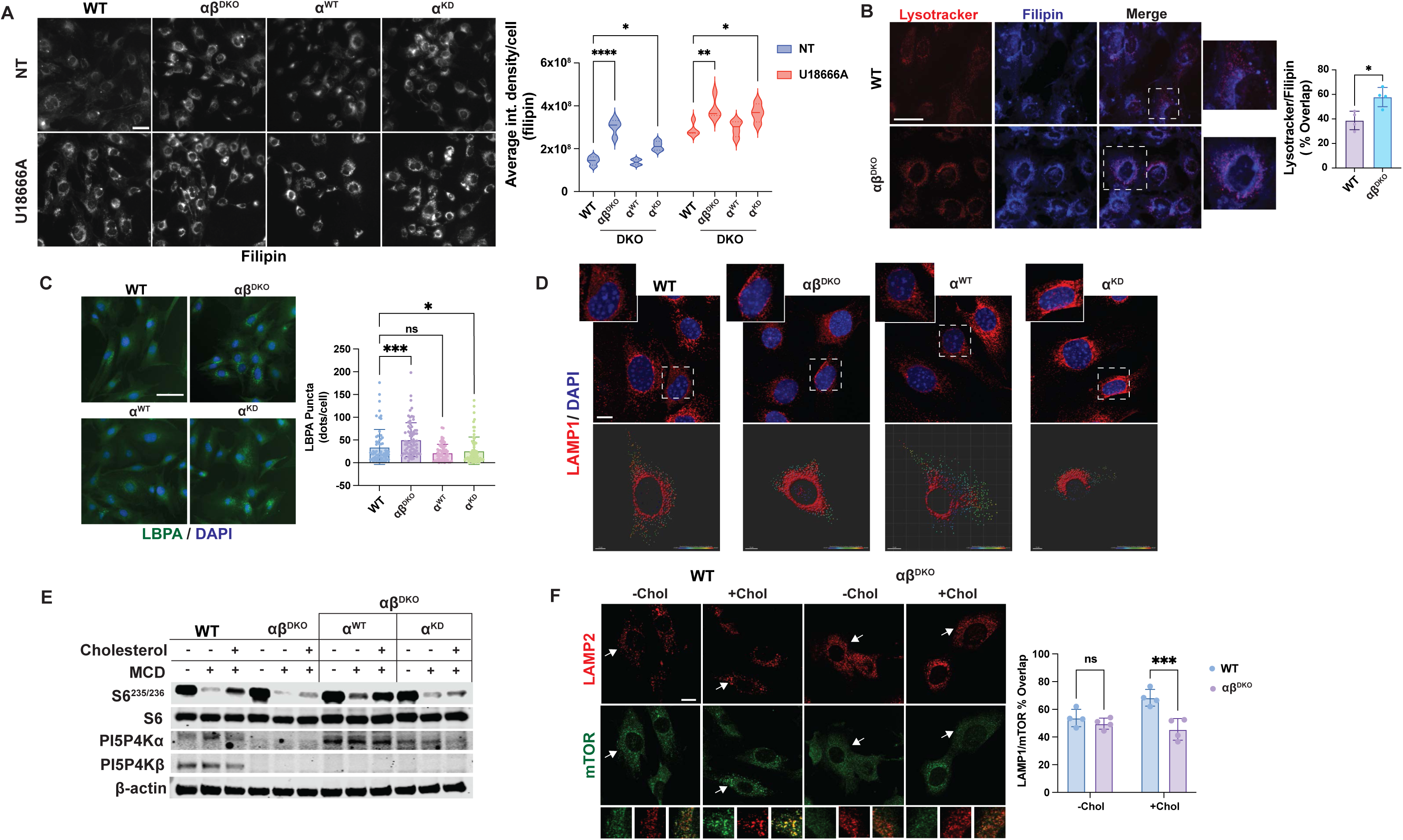
PI5P4Ks support cholesterol homeostasis through lysosome positioning and mTOR activation. **(A)** Free cholesterol staining of Trp53-/- MEFs with filipin. Cells were either non-treated (NT=10% FBS) or treated with an NPC1 inhibitor for 16 hrs (U18666A) to induce cholesterol accumulation. WT=*Pip4k2a+/+ Pip4k2b+/+*, abDKO=*Pip4k2a-/- Pip4k2b-/-*, aWT=functional PI5P4Ka rescue, aKD=PI5P4Ka kinase dead. Scale bar=50uM. Shown are mean+SD. ANOVA followed by Dunnett’s multiple comparison correction, compared to WT. p=*0.05, **0.01, ***0.001, ****0.0001. **(B)** Filipin and lysotracker overlap in untreated *Trp53-/-* MEFs. Scale bar=50uM. Shown is mean+SD. Student’s t-test. p=*0.032 **(C)** LBPA staining in untreated *Trp53-/-* MEFs. Shown is mean+SD obtained from CellProfiler analysis. ANOVA followed by Dunnett’s multiple comparison correction, compared to WT. p=*0.05, ***0.001. **(D)** Lysosome positioning analysis of Trp53-/- MEFs using LAMP1 as a lysosomal marker. Images were acquired with confocal microscopy at 63x magnification (oil) and representative of 5 cross-sections of 0.5uM Z-stacks with maximum intensity projection. Scale bar=10uM. **(E)** Immunoblot of Trp53-/- MEFs for PI5P4Ks and mTORC1 target S6 in a cholesterol depletion and repletion assay. Sterol depletion was performed using methyl-β-cyclodextrin (MCD, 1% w/v) for 1.5 hours, followed by refeeding for 1.5 hours with 150 μM cholesterol. **(F)** Immunofluorescence of endogenous mTOR and LAMP2 colocalization in Trp53-/- MEFs. Cholesterol depletion with MCD for 1.5 hr (-Chol) compared to depletion + readdition of cholesterol for 1.5 hr each (+Chol). Images were acquired with confocal microscopy at 63x magnification (oil). Scale bar=10uM. Shown is the mean+SD of the calculated overlap. Unpaired t-test followed by Holm-Sidak correction. p=**0.0037.

Based on these results, we sought to determine the consequence of diminished cholesterol efflux to other cellular compartments upon loss of PI5P4K function. In NPC1-deficient cells, failed cholesterol efflux can cause an expansion of cholesterol-laden lysosomes and endosomes, which contain lysobisphosphatidic acid (LBPA) on the membrane (*36, 37*). Using a monoclonal LBPA antibody, the p53-/- aβ^DKO^ and p53-/- PI5P4K⍺^KD^ conditions both had a substantial increase in LBPA staining in comparison to the p53-/- WT and p53-/- PI5P4K⍺^WT^ counterparts (Fig. 2C).

This expansion of cholesterol-containing endosomes in PI5P4K deficient cells suggests a downstream effect on endosomal trafficking (*38*). Therefore, we investigated the consequence of changes in lysosomal positioning as an affected process of cells lacking PI5P4Ks. Lysosomal positioning is a well-studied phenomenon in which nutrient cues in the cell influence the dynamic movement of lysosomes along microtubules to either the periphery or the perinuclear area of the cell. Therefore, we examined the distribution of where these cholesterol-enriched lysosomes were positioned intracellularly. The positioning of lysosomes and late endosomes has been associated closely with the mTORC1 signaling pathway, whereas the localization of mTORC1 to the lysosomal membrane is critical for mTORC1 activation (*39, 40*). In the nutrient-sufficient condition, lysosomes are positioned toward the cell periphery, where they are detected at a higher coincidence with mTORC1 due to the proximity of the cell surface and signaling receptors (*41*). During states of nutrient depletion, lysosomes are found at a higher ratio in the perinuclear area, where they are less likely to interact with mTORC1 and are primed for autophagy and fusion events to protect cells from starvation. We found that in the p53-/- αβ^DKO^ and p53-/- PI5P4K⍺^KD^ conditions, lysosomes were accumulated towards the perinuclear region in comparison to p53-/- WT and p53-/- PI5P4K⍺^WT^ cells (Fig. 2D).

As the perinuclear clustering of lysosomes should, in turn, be concurrent with mTORC1 activity, we investigated downstream activation of the mTOR pathway via phosphorylation of S6 protein at serine 235/236 (pS6^S235/236^). Cholesterol-dependent mTOR signaling was investigated using a cholesterol depletion and repletion assay, in which cholesterol is depleted using methyl-β-cyclodextrin (MCD) treatment (1% w/v) followed by cholesterol repletion (50uM) for 1.5 hours. Cells lacking PI5P4Ks failed to recover from cholesterol depletion and showed minimal phosphorylation of pS6^S235/236^. In contrast, p53-/- WT and p53-/- PI5P4K⍺^WT^ cells responded almost to baseline activation levels (Fig. 2E). We reasoned that this deficiency in mTORC1 activation could be due to altered lysosome positioning and decreased mTOR localization to the lysosome. Therefore, we performed immunofluorescence on these cell lines to quantify the overlap of mTOR and the lysosomal marker LAMP1. We found that in conditions where mTORC1 signaling was decreased, there was a concurrent decrease in mTORC1 localization at the lysosome in cells lacking PI5P4Ks (Fig. 2F).

These data indicate that cells lacking PI5P4Ks are prone to cholesterol accumulation due to lack of PI(4,5)P_2_, as PI5P4K⍺^WT^ exhibited a rescue of the phenotype, but this was not observed in the PI5P4K⍺^KD^ condition. Overall, we find that the activity of the PI5P4Ks is necessary for the export of lysosomal cholesterol in the p53-/- context. Further, the lack of PI5P4K activity increases the pool of late endosomes, including lysosomes positioned in the perinuclear area. This leads to a reduction in mTORC1 association with lysosomes and disrupts lysosomal nutrient sensing, with downstream consequences on autophagy and activation of mTOR.

### Deletion of PI5P4Ks in p53-deficient human cells dysregulates lysosome positioning and mTOR growth pathway

It was recently discovered that the oxysterol binding protein (OSBP), typically found at contact sites between the ER and trans-Golgi network, was clustered on LAMP2-positive lysosomes in HEK29A cells (*21*). The OSBP-related proteins (ORPs) contain large hydrophobic domains at their C-terminus, which bind and protect hydrophobic molecules from the cytosol, including cholesterol and phospholipids (*42*). The localization of ORPs at the lysosomes of these cells provided our rationale for the use of HEK293A as a suitable model to examine lysosome cholesterol trafficking (*21*). To examine the loss of PI5P4Ks in this system, we performed CRISPR-Cas9 mediated deletion of *PIP4K2A* and *PIP4K2B* to generate (PI5P4K) αβ^DKO^ cells. We then introduced a stable shRNA knockdown of *TP53* (shp53) in HEK293A αβ^DKO^ and WT (control) cells to model the dysregulation of cholesterol biosynthesis as a result of p53 loss.

To measure specific pools of cholesterol at membranes, we utilized a biosensor (eOsh4-WCR) derived from the engineered sterol transport protein, Osh4, conjugated to an environment-sensitive amphiphilic fluorophore, WCR (*43, 44*). Spatiotemporal measurement of lysosomal cholesterol concentration was achieved by co-transfecting the cells with the biosensor and eGFP-LAMP1 to measure lysosome-specific cholesterol signals and to normalize lysosome counts between conditions. Notably, the spatially averaged cholesterol concentration at the cytofacial plasma membrane was approximately equal in both conditions. However, the spatially averaged cytofacial lysosome cholesterol concentration was increased 1.53 fold in shp53 αβ^DKO^ compared to shp53 WT control (Fig 3A).

**Figure 3.**
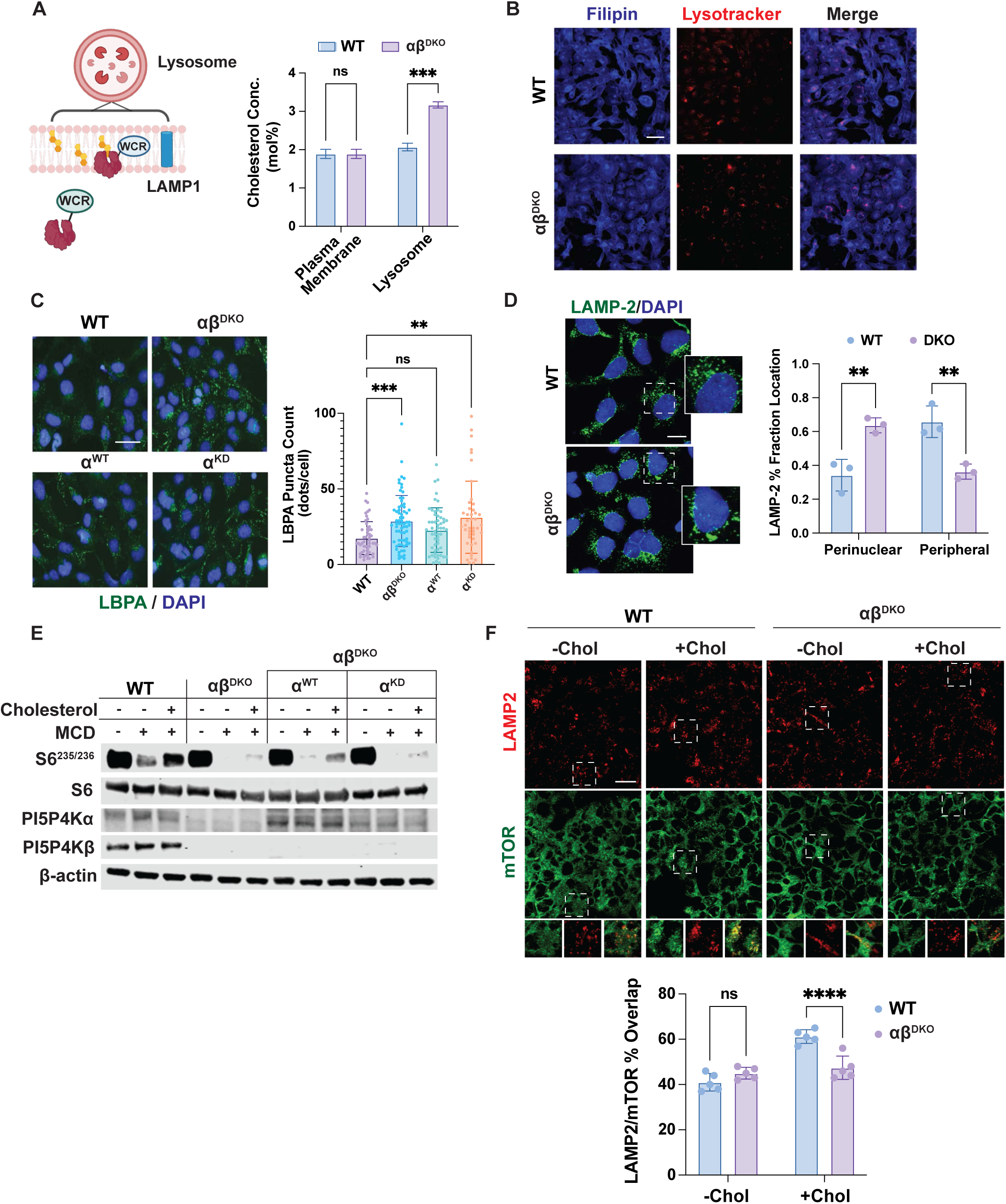
Deletion of PI5P4Ks in p53-deficient cells dysregulates lysosomal cholesterol handling and perturbs mTOR growth pathway. **(A)** Ratiometric determination of membrane cholesterol levels in 293A shp53 cells using cholesterol sensing domain Osh4 conjugated to a solvatochromic fluorophore (WCR). Lysosomal measurement quantified by overlap with LAMP1 marker and normalized per lysosome count. N=2 independent replicates. Unpaired t-test mean+SD. p=***0.0001. **(B)** Visualization of late endosomes/lysosomes in 293A shp53 cells using filipin and lysotracker. Scale bar=200uM. **(C)** 293A shp53 cells fixed and stained with antibody to LBPA and imaged at 20x magnification. Images are representative of three independent replicates. Statistics represent two-way ANOVA with multiple comparisons compared with control (WT). Dunnett’s test used for multiple comparison correction, p-value <0.05*, 0.01**, 0.001***.**(D)** 293A shp53 cells stained for lysosomal marker LAMP-2 for analysis of lysosome positioning. N=3 independent replicates. Shown is mean+SD. ANOVA followed by Sidak correction for multiple comparisons. **p=0.002. **(E)** Immunoblot of 293A shp53 cells for PI5P4Ks and mTORC1 target genes in cholesterol depletion and repletion assay. **(F)** 293A shp53 cells costained for endogenous mTOR and LAMP-2. Scale bar=10uM. Shown is the mean+SD of the calculated overlap. ANOVA with Bonferroni correction for multiple comparisons. ns=0.2678, p=****<0.0001

We next performed filipin staining with lysotracker to visualize the increase in cellular free cholesterol and found a significant increase in shp53 αβ^DKO^ (Fig 3B). Interestingly, in the shp53 αβ^DKO^, we observed an increase in lysotracker staining compared to the shp53 WT, which could be attributed to a decreased lysosomal turnover rate due to failed autophagosome-lysosome fusion (Fig. S3A) (*14*).

To verify this accumulation of lysosomes and cholesterol was due to the loss of PI5P4K activity rather than deletion of the protein, we reintroduced the functional PI5P4Kα (α^WT^), as well as the kinase-dead PI5P4Kα (α^KD^) into the shp53 αβ^DKO^ cells and performed staining of LBPA, which is found primarily in cholesterol-enriched intraluminal vesicles of the late endosome/lysosome compartment (*45, 46*). We found that shp53 WT and α^WT^ cells had a significant decrease in LBPA staining compared to both the shp53 αβ^DKO^ and α^KD^, suggesting that the catalytic function of PI(4,5)P_2_ generation was sufficient to rescue endolysosomal cholesterol accumulation (Fig. 3C). Of note, we observed that LBPA staining was enhanced at the perinuclear compartment in shp53 αβ^DKO^ and α^KD^, suggesting that cholesterol enrichment of these compartments was impacting late endosome/lysosome positioning.

Lysosome positioning has been shown to modulate both activation of the mTORC1 growth signaling pathway and autophagy (*40*). To investigate changes in lysosomal positioning upon loss of the PI5P4Ks, we performed staining of the lysosomal marker LAMP2. Similar to the MEF system, we found that cells lacking PI5P4Ks exhibited a significant shift to the perinuclear compartment (Fig. 3D). To examine if the cholesterol-dependent reactivation of downstream mTORC1 targets was consistent from murine to human cells, we performed acute-starvation and refeeding experiments with cholesterol over a time course of 60 minutes or for 1.5 hours, respectively (Fig. S3B, Fig. 3E). In the shp53 WT cells, cholesterol depletion leads to a decrease in mTOR pathway activation observed by a decrease in pS6^235/236^ signal (Fig. 3E). Interestingly, this depends on kinase activity, as shp53 α^WT^ can rescue mTOR activation but not shp53 α^KD^ (Fig 3E). In addition, we observed a significantly decreased colocalization between mTOR and the lysosome marker, LAMP2, upon cholesterol repletion in shp53 αβ^DKO^, while cells observed in the fed condition (10% FBS) only had a minor decrease (Fig 3F, S3C). These findings suggest the PI5P4Ks are necessary for cholesterol clearance from the lysosome and that the lack of clearance in the absence of PI5P4Ks results in the mislocalization of lysosomes to the perinuclear area. This mislocalization of lysosomes, in turn, leads to the diminished capacity for lysosome colocalization with mTORC1, preventing downstream activation of this growth pathway for cellular growth signaling, even in cells lacking p53.

### Breast cancer cells require PI5P4Ks for survival and cholesterol sensing

In breast cancers, and especially triple-negative breast cancers, the PI3K/Akt/mTOR (PAM) pathway is one of the most active pathways involved in survival and chemoresistance (*24, 25*). To further test the hypothesis that PI5P4Ks are required in the p53-deficient context to maintain lysosome positioning and mTORC1 activation, we utilized a panel of human breast cancer cell lines. We performed shRNA depletion of both PI5P4K isoforms to generate PI5P4K αβ^KD^ cells to compare with PI5P4K non-targeting scramble (Scr). As dual inhibition of PI5P4Kα and PI5P4Kβ significantly affects cell proliferation, we generated PI5P4K αβ^KD^ in two steps. First, we generated stable shRNA PI5P4Kβ knockdown cells, followed by stable shRNA PI5P4Kα knockdown. Two independent shPI5P4Kα hairpins were used, αβ^KD#1^ and αβ^KD#2^. In MCF-7 cells, which express wild-type p53, no significant inhibition of growth was observed upon PI5P4K knockdown (Fig. 4A). In contrast, cells of p53-deficient genetic background had a significant proliferation defect upon knockdown (Fig. 4A, S4A). To investigate transcriptional changes upon PI5P4K knockdown in cancer cells, we performed RNA-sequencing in the p53-deficient, TNBC cell line, HCC1806. Importantly, we validated that the two independent hairpins were inducing similar transcriptional changes in these cells (Fig. S4B). Gene set enrichment analysis (GSEA) revealed mTORC1 pathway downregulation with concurrent upregulation of lysosomal genes (Fig. 4B, S4B). In addition, HCC1806 were responsive to the cholesterol depletion and repletion assay, whereby αβ^KD^ impaired cholesterol-induced mTORC1 activation, evident by immunoblot of the targets pS6^S235/236^ and p4EBP1^S65^ (Fig. 4C). To investigate these findings in an *in vivo* setting, we introduced a doxycycline-inducible shRNA to PI5P4Kα in the stable knockdown PI5P4Kβ cells (HCC1806 PI5P4K αβ^in-KD#1^ and PI5P4Kαβ^in-KD#2^) to compare with control cells containing an inducible scramble shRNA (HCC1806 Scr^in^) to circumvent the lethality caused by stable knockdown (Fig. 4D). To preserve the tumor context, these cells were then orthotopically implanted into the mammary fat pads of immune-deficient Fox1*Nu/Nu* mice. Once tumors reached an average volume of 80mm^3^, mice were treated with doxycycline to induce the silencing of PI5P4Kα (Fig. 4D, Fig. S4D). Upon knockdown, HCC1806 PI5P4Kαβ^in-KD^ tumors were severely growth impaired over a 14-day growth period (Fig. 4E, Fig. S4E). To determine if mTORC1 signaling was impaired as a consequence of PI5P4K knockdown *in vivo*, tumor immunostaining was performed for downstream phosphorylation targets, pS6^S235/236^ and p4EBP1^S65^. A significant decrease in staining was detected in the tumors lacking PI5P4Ks, suggesting they are required for mTORC1 activation in the tumor (Fig. 4F). To expand these findings to cholesterol-induced activation based on lysosomal cholesterol transport, we also stained these tumors for filipin and the lysosome marker LAMP2 and found that tumors lacking the PI5P4Kα and PI5P4Kβ isoforms had increased intratumoral cholesterol levels that coincide with the lysosomal compartment (Fig. 4G). We determined this growth impairment observed upon PI5P4K knockdown was a result of cell death, rather than an impairment of proliferation, based on enhanced staining of the apoptotic marker, cleaved-caspase-3, and no change in the mitotic marker, phospho-histone H3 (Fig. S4F). Taken together, these results indicate that PI5P4Ks are necessary for proliferation, cholesterol trafficking, and maintaining mTORC1 signaling in p53-deficient TNBC cells, both *in vitro* and *in vivo*.

**Figure 4.**
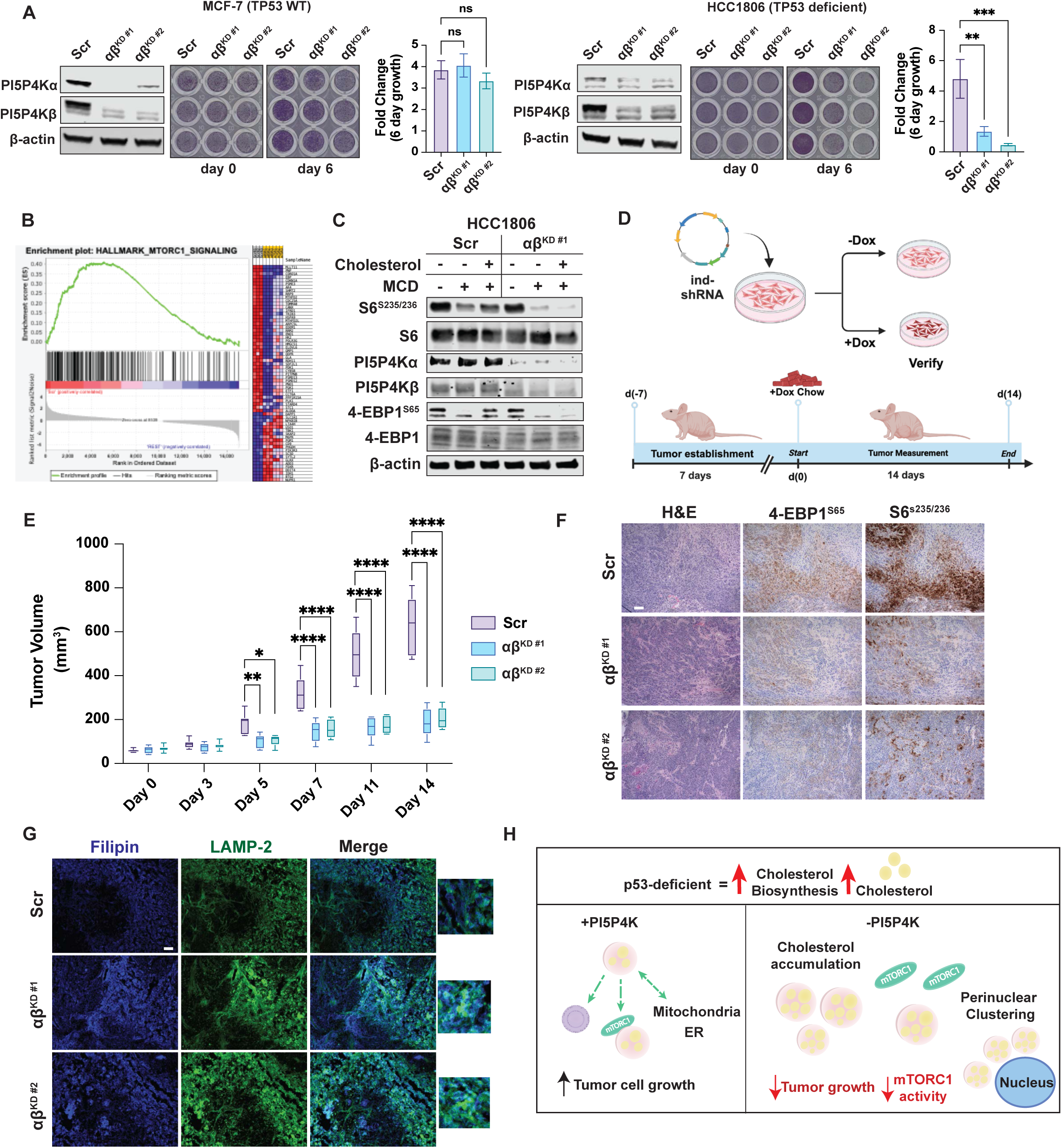
Breast cancer cells require PI5P4Ks for survival and cholesterol sensing. **(A)** Proliferation assays were performed using crystal violet on indicated cell lines with either non-targeting scramble shRNA (Scr) or dual, stable shRNA knockdown of PI5P4Ks (αβKD#1 or αβKD#2). N=3 independent replicates. Shown is mean + SD. ANOVA followed by Dunnett’s multiple comparison correction compared to Scr. p=*0.01, **0.001, ***0.0001. **(B)** RNA-sequencing of p53-deficient HCC1806 cells was analyzed for pathway analysis using GSEA. Signatures for Hallmarks mTORC1 signaling (NES=(+)2.0, p=0.0). **(C)** Immunoblot of HCC1806 cells (Scr vs αβKD#1) in a cholesterol depletion and repletion assay. **(D)** Schematic of HCC1806 in vivo orthotopic breast implantation development and timeline. **(E)** HCC1806 tumor progression over time comparison of inducible hairpins; Scr, αβKD#1, and αβKD#2. Treatment with doxycycline began when tumors reached 75mm3 (Day 0). Shown is a bar plot of mean+SD. ANOVA with multiple comparisons to Scr with Sidak correction. p<*0.01, ***0.001, ****0.0001. **(F)** Immunohistochemistry of HCC1806 tumors stained for H&E and the mTORC1 targets p4EBP1^S65^ and pS6^S235/236^. Scale bar=100uM. **(G)** Immunofluorescence staining of HCC1806 tumor slices stained for filipin and LAMP2 to visualize cholesterol overlap. Scale bar=100uM. **(H)** Model for PI5P4K involvement in lysosomal cholesterol handling and lysosome localization in p53-deficient cells.

## Discussion

Investigating the role of the PI5P4Ks at the lysosome has great potential for understanding the biology and the pathophysiology of lysosome-related dependencies in cancer and other metabolic diseases. Although these enzymes are often difficult to implicate in cancer using bioinformatics due to their lack of mutations, we have demonstrated that their elevated expression does correlate with several critical metabolic nodes that have been investigated in this work, including cholesterol trafficking and mTOR signaling. Our findings reveal, for the first time, that the complete loss of PI5P4Kα and PI5P4Kβ alleles in adult mice is non-lethal and confers protection from tumorigenesis in the p53-deficient mouse context. Previously, we have attributed this synthetic lethal phenotype to the notion that the PI5P4Ks have evolved as critical modulators of the cellular stress response, with roles in protection from oxidative stress, mitochondrial health, and modulators of autophagy (*13, 14, 47*). We have now identified a previously undescribed role for the PI5P4Ks in maintaining lysosomal cholesterol homeostasis and mTORC1 signaling. These findings, substantiated in MEFs and HEK293A cells, suggest a new upstream rationale for the previously discovered autophagy and mitochondrial defects in p53-deficient cells lacking PI5P4Ks (*14, 48*).

An interesting notion of this research is that the accumulation of lysosomal cholesterol causes a buildup of lysosomes in the perinuclear area and a decrease in cholesterol-mediated mTORC1 localization to the lysosomal membrane. We determine that the PI5P4Ks are necessary for cholesterol clearance from the lysosome and that the lack of clearance in the absence of PI5P4K-mediated PI(4,5)P_2_ generation results in the mislocalization of lysosomes to the perinuclear region. Mislocalized lysosomes lead to the diminished capacity for lysosome colocalization with mTORC1, preventing downstream growth pathway activation. Indeed, this finding is paradoxical, as patients harboring mutations in the lysosomal cholesterol export protein NPC1 typically exhibit mTOR hyperactivation (*20*). We currently do not have a rationale for why this occurs, although several groups have reported an impairment in mTORC1 activation upon inhibition or loss of PI5P4Ks, which could potentially be the result of an impairment in the autophagic response and lysosome-driven metabolic processes.

Using this rationale, we determine that the PI5P4Ks are necessary for cholesterol clearance from the lysosome and that the lack of clearance in the absence of PI5P4K-mediated PI(4,5)P_2_ generation results in mislocalization of lysosomes to the perinuclear region. Mislocalized lysosomes lead to the diminished capacity for lysosome colocalization with mTORC1, preventing downstream growth pathway activation (Fig. 4H).

Besides uncovering a novel role for the PI5P4Ks in coordinating lysosomal cholesterol trafficking, this study demonstrates for the first time the critical nature of these enzymes in triple-negative breast cancer tumor maintenance. Using an inducible knockdown model, we have shown that depletion of these enzymes in established tumors leads to decreased activation of mTORC1 signaling, apoptotic cell death, and an accumulation in intratumoral cholesterol trafficking at the lysosome. An intriguing question that remains is to what extent the deficiency of p53 impacts endogenously synthesized cholesterol turnover, LDL-derived cholesterol import, or the balance between these two systems. This question will have great importance for the future of therapeutics where cholesterol trafficking is targeted in breast cancer and beyond.

## Materials and Methods

### Cell Lines

All cells were incubated in a 37C humidified incubator at 5% CO2. MEFs, HEK293A, HEK293T, MCF-7, MDA-MB-436, MDA-MB-361, and HCC1806 cells were cultured in Dulbecco’s Modified Eagle Medium (DMEM Corning 10-013-CV) supplemented with 10% Fetal Bovine Serum (FBS), penicillin/streptomycin (100 U/mL). HEK293T, HEK293A, MCF-7, MDA-MB-436, MDA-MB-361, and HCC1806 were obtained from American Type Culture Collection (ATCC). MEFs were isolated from E13.5 embryos from the following lines: *Pip4k2afl/fl Pip4k2b-/- Trp53fl/fl* and *Pip4k2a+/+ Pip4k2b+/+ Trp53fl/fl*. After two passages, MEFs were treated with 4-Hydroxytamoxifen in culture for five consecutive days (4-OHT; 10ug/mL) to induce recombination of floxed alleles *in vitro*. Successful recombination was confirmed by genotyping. MEF DNA was prepared from the head of embryos using a Qiagen DNeasy kit and genotyped for Pip4k2a and Pip4k2b using the primer pairs as previously described (*13*). *Trp53* was genotyped using the following primer set for the Berns alleles recombined from the Jacks Lab website: (https://jacks-lab.mit.edu/protocols/genotyping/). MEFs used for experiments had the following genotypes: *Pip4k2a-/- Pip4k2b-/- Trp53-/-* and *Pip4k2a+/+ Pip4k2b-/- Trp53-/-*.

Cell line authentication was performed (prior to freezing initial early-passage stocks) on HEK293A, MCF-7, MDA-MB-436, MDA-MB-361, and HCC1806 cell lines by the Genomics core at Sanford Burnham Prebys using STR analysis. Mycoplasma testing was performed using the ABM Mycoplasma PCR detection kit upon arrival of cell lines and monthly during culture (ABM, G238). Early-passage cells of parental lines were frozen and maintained no longer than ten passages at a time.

### Viral Transduction of Cell Lines

The 293T packaging cell line was used for lentiviral amplification. Briefly, viruses were collected 48 hours after transfection, filtered, and used for infecting cells in the presence of polybrene (10 μg/ml) before puromycin selection. pVSVg and pPax were used for lentiviral packaging.

Stable knockdown of *PIP4K2A* and *PIP4K2B* was achieved using lentiviral transduction of shRNA. HEK293T cells were cotransfected with 12ug of each shRNA containing pLKO.1 plasmid, 8ug of psPAX2 packaging plasmid, and 4ug of pMD2.G envelope plasmid using Lipofectamine 2000 in 10cm dishes. After 24 hours, the medium was replaced with fresh 10% FBS DMEM without penicillin/streptomycin. After 48 hours, viral supernatant was collected and filtered using 0.45uM cellulose acetate syringe filters. Cancer cells were transduced with viral supernatant and polybrene (8ug/mL) for 16 hours. Due to the lethality of dual knockdown in cancer cell lines, knockdowns were performed successively, whereas *PIP4K2B* stable knockdown cells were transduced first. Following transduction with *PIP4K2B* viral supernatant, selection was achieved using Geneticin (0.5-2mg/mL) for two weeks. The knockdown of PIP4K2B was confirmed by western blot, and early passage cell stocks were frozen. Next, transduction with PIP4K2A viral supernatant was performed, and selection was achieved using puromycin (0.5-2ug/mL) for two days. Post-selection cells were split for experiments and simultaneous confirmation of knockdown. Inducible knockdown of *PIP4K2A* was achieved using the lentiviral pLKO.1 system in a similar manner. HCC1806 inducible knockdown cells were first generated with stable *PIP4K2B* knockdown and selection with Geneticin (0.5-2mg/mL) and followed by *PIP4K2A*-inducible transduction and selection with puromycin. To avoid doxycycline-inducible hairpin activation, PIP4K2A-inducible stable cells were maintained in Tet system-approved FBS.

### Immunoblot analysis and antibodies

Total cell lysates were prepared by washing cells with cold phosphate-buffered saline. The cells were then lysed with buffer containing 20 mM Tris/HCl (pH 7.5), 150 mM NaCl, 1 mM EDTA, 1mM EGTA, and 1% Triton, with the addition of protease and phosphatase inhibitors. Protein was measured using the BCA assay, and 20-35 μg of total cell lysates were run on SDS–polyacrylamide gel electrophoresis. The proteins were transferred onto a nitrocellulose membrane, and membranes were probed overnight at 4°C with the appropriate primary antibody. Antibodies used were as follows: PIP4K2α (5527; Cell Signaling), PIP4K2β (9694; Cell Signaling), P-p70S6K (9234; Cell Signaling), p70S6K (9202; Cell Signaling), P-S6 (2211; Cell Signaling), S6 (2217; Cell Signaling), P-4EBP1 (2855; Cell Signaling), 4EBP1 (9452; Cell Signaling), p53 (2524; Cell Signaling), α-tubulin (T6199; Sigma), and β-actin (ab8226; Abcam).

### Immunofluorescence

Cells were plated on either acid-washed glass coverslips or 96-well optical-grade plastic plates. Post-treatment, cells were fixed using methanol-free paraformaldehyde (4% PFA) diluted in PBS for 15 minutes at room temperature. Fixative was quenched with 15mM glycine/PBS with two 5-minute washes. Cells were then permeabilized with 0.1% saponin/PBS for 10 minutes. Post-permeabilization cells were blocked with 1% BSA/0.01% saponin/PBS for 30 minutes. Primary antibodies were diluted in the same blocking buffer and added for 1 hour at room temperature or overnight at 4°C. Cells were washed three times with blocking buffer for 5 minutes each, and fluorescent secondary antibodies were added at 1:1000 for 1 hour at room temperature. Cells were washed again three times and either mounted in DAPI containing mounting media or DAPI was added in an additional blocking buffer wash for 10 minutes to stain the nuclear compartment. Slides or plates were imaged immediately or stored at 4°C in the dark.

### Cholesterol manipulation of cells

p53-/- MEFs, shp53 HEK293A, or HCC1806 cells in culture dishes were rinsed once with serum-free media and incubated in DMEM containing 1.0% methyl-β-cyclodextrin (MCD) (Sigma C4555) supplemented with 0.5% lipid-depleted serum (LDS) for 1.5 hours. Cells were then transferred to DMEM supplemented with 0.1% MCD and 0.5% LDS (-Chol) or to DMEM containing 50 uM cholesterol complexed to MCD (Sigma C4951) for solubility and incubated for an additional 1.5 hours.

### RNA-sequencing and analysis

HCC1806 cells with shRNA knockdown of PI5P4Ks were performed four days before collection for RNA sequencing. Total RNA was prepared using Direct-zol RNA MiniPrep on ice and stored at -80°C before analysis. RNA yield and purity were tested using the Qubit 4 Fluorometer before analysis. Sequencing libraries were prepared using 250 ng of RNA using standard Illumina TruSeq single indexing protocols and sequenced using the Illumina NextSeq 500 instrument. Adapter remnants of sequencing reads were removed using cutadapt v1.18. Read alignment was performed using human genome v.38 and Ensembl gene annotation v.84 using STAR aligner v.2.7. DESeq2 was used for differential gene expression analysis. RNA sequencing results have been deposited in GEO under accession number GSE221217.

### Gene set enrichment

GSEA was performed using GSEA v4.0.3. Normalized enrichment scores (NESs) and P values were used to determine the significance of the findings.

### Analysis of lysosome positioning

Lysosome positioning was analyzed using a custom CellProfiler pipeline. In brief, the identification of primary object modules was used to identify nuclei, LAMP1/2 spot detection, and segment cell borders. Masks were then generated to extend 10um from the nuclear region to identify the perinuclear region. Next, a 5um spacer region was identified to add distance between perinuclear and peripheral regions. Lastly, a peripheral region encompassing the outer perimeter space away from the perinuclear region was defined. LAMP1 signal was calculated for both the perinuclear and peripheral regions and calculated into a ratio.

### Cholesterol biosensor preparation of WCR-eOsh4

The eOsh4 was transformed to E. coli BL21 RIL codon plus cells (Stratagene) for bacterial expression. A preculture was prepared from a single colony in 10 ml of Luria-Bertani medium with 50 μg/ml kanamycin and was incubated in a shaker overnight at 37°C or until it got cloudy. Ten milliliters of preculture were transferred to 1 l of main culture with 50 μg/ml kanamycin and incubated in a shaker at 37°C until A600 reached 0.6. Then, protein expression was induced at 18°C with 0.5 mM isopropyl β-d-1-thiogalactopyranoside for 16 h. The induced culture was aliquoted to 250 ml and centrifuged at 4,000 g for 10 min. Cell pellets were stored at −80°C until use. The cell pellets were resuspended with 20 ml of the lysis buffer (50mM Tris-HCl [pH 7.9], 300 mM NaCl, 10 mM imidazole, 10% glycerol, 1 mM phenylmethanesulfonylfluoride, and 1mM dithiothreitol) and lysed by sonication. The lysate was centrifuged at 44,000 g for 30min and the supernatant was mixed with 1 ml of Ni NTA agarose resin (Marvelgent Biosciences Inc.) and incubated at 4°C for 2 h with gentle shaking. For preparation of WCR-eOsh4, the eOsh4-bound resin was resuspended with WCR (1:10 molar ratio) in 5 ml of labeling buffer [50 mM Tris, pH 8.05, containing 150 mM NaCl, 20 mM imidazole, 50 mM arginine, 50 mM glutamate, and 1 mM Tris(2-carboxyethyl)phosphine (TCEP)] and the mixture was gently shaken for 2 h at room temperature, or at 4°C overnight in a gyratory shaker. WCR-eOsh4 was then washed with 50 ml of the wash buffer (80 mM Tris, pH 7.9, 300 mM NaCl, 40 mM imidazole) containing 4% (v/v) dimethyl sulfoxide and then with 300 ml of the wash buffer. WCR-eOsh4 was eluted from the resin with the elution buffer (50 mM Tris, pH 7.9, 300 mM NaCl, 300 mM imidazole). Collected fractions were concentrated in an Amicon Ultra 0.5 ml Centrifugal Filter (Millipore), and the buffer solution was exchanged to 20 mM Tris, pH 7.4, 160 mM NaCl. The protein concentration of the WCR-eOsh4 solution was determined by the Bradford assay. All steps were performed at 4°C unless otherwise mentioned.

### Cholesterol biosensor in situ quantitative imaging

The same number (2.5 × 10^4^) of cells were seeded into 100 mm round glass-bottom plates (MatTek) and grown at 37°C in a humidified atmosphere of 5% CO2 in DMEM supplemented with 10% (v/v) FBS, 100 U/ml penicillin G, and 100 mg/ml streptomycin sulfate and cultured in the plates for about 24 h and transfected with LAMP1 as lysosome marker using JetPRIME system overnight before lipid quantification. Imaging was performed with the custom-designed six-channel Olympus FV3000 confocal microscope with the environmentally controlled full enclosure incubator (CellVivo). Cells were maintained at 37°C and with 5% CO2 atmosphere throughout the imaging period to maintain the cell viability. Typically, 20–30 fl of the sensor solution was microinjected into the cell to reach the final cellular concentration of 200–400 nM. All image acquisition and imaging data analysis, as well as the GUV calibration curves, are done with Image-proPlus 7, as described previously (*43*).

### Human cancer bioinformatics

Bioinformatics analysis of human data was performed using cBioPortal. GSVA analysis of the TCGA PanCancer BRCA dataset was performed in R. Selected pathways from MSigDB were used for PIP4K2A quartile analysis. cBioPortal was used to subset TCGA BRCA patients into four clinical subtypes based on clinical expression data of markers. For paired vs normal tumor analysis, TNMplot was used for gene expression analysis (*49*).

### Orthotopic implantation of human breast cancer cells

Inducible non-targeting scramble or PI5P4K knockdown HCC1806 cells were maintained in Tet-free FBS before implantation. Cells were trypsinized and resuspended in PBS to obtain a suspension of 5 * 10 ^ 5 cells per injection, then mixed into a 1:1 suspension of matrigel/PBS. Cell suspensions were kept on ice at all times prior to injection to prevent solidification of matrigel. *Fox1^Nu/Nu^* mice were received from Jackson Labs and acclimated to the procedure room for 2 weeks prior to experimental injections. Mice were anesthetized on a heated stage using isofluorane, and toe pinch was used before switching mice to nosecone to ensure proper anesthetic depth. The surgical site was sterilized using three consecutive wipes of betadine and ethanol and all tools were autoclaved prior to the surgical procedure. A small incision (∼3mm) was made 5mm beneath the nipple of the 4th and 9th inguinal fat pads using operating scissors. Using the blunt edge of the scissors, the skin inside the incision was loosened from the body wall until the mammary fat pad could be visualized. Using forceps, the fat pad is gently guided through the incision, and 100uL of cell suspension is delivered directly to the fat pad. The fat pad is then gently placed back into position, and the incision is closed using surgical glue. Mouse recovery is monitored post-procedure to ensure closure of the surgical site and full recovery of motion. Mice were then monitored for tumor growth every two days until tumors reached ∼80mm^3^, measured with digital calipers. Once initial tumor size was reached, mice were placed on a 625 mg/kg doxycycline chow diet for the duration of the experiment to induce PI5P4K knockdown.

### Statistics and analysis software

Data are expressed as means ± SD unless otherwise specified. Data were verified for normality using the Shapiro-Wilk normality test, and data failing normality were analyzed using non-parametric tests. Statistical analyses for all data are indicated in the figure legend and were performed using GraphPad Prism 10 (software version 10.3.0). Statistical significance is indicated in the figures. *p < 0.05, **p < 0.01, ***p < 0.001, ****p < 0.0001, unless specified otherwise.

## Acknowledgments

We would like to thank the following cores at SBPMDI (NCIP30CA030199): Cell Imaging and Histology, Bioinformatics, and Genomics. We would also like to thank the Animal Facility and IACUC board members at SBPMDI for assisting in the care and use of our laboratory animals. We thank C. Commisso, G. Wahl, P. Adams, and M. D’Angelo for their insightful discussions and their input on the project.

## Author contributions

Conceptualization: RML, GKA, ALL, BME, JS, WC. Methodology: RML, GKA, ALL, BME, SC, KL, RH, JS, WC. Investigation: RML, SC, KL, RH, JS, BME, WC. Visualization: RML, BME. Funding acquisition: RML, BME, WC. Project administration: RML. Supervision: BME. Writing – original draft: RML, BME. Writing – review & editing: RML, BME, GKA, ALL, SC

## Data and materials availability

All data are available in the manuscript, the supplementary material, or the Gene Expression Omnibus (Accession # GSE221217), which contains the processed RNA-seq for cell line HCC1806. All materials are available from the corresponding author upon reasonable request.

---

**Figure S1.**
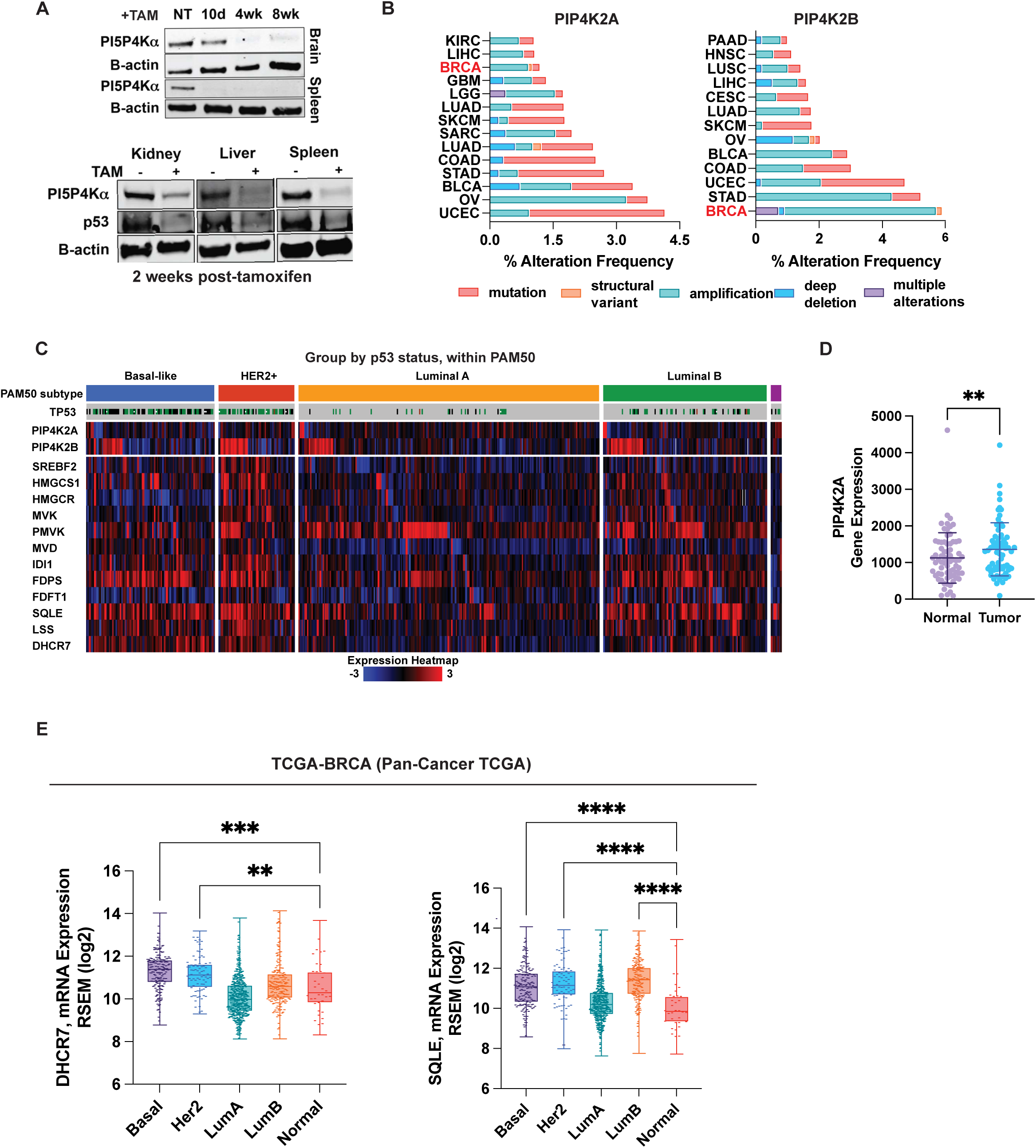
**(A)** Validation of protein knockout post-tamoxifen treatment at labeled time points and at two weeks post tamoxifen induction. **(B)** Distribution of *PIP4K2A* and *PIP4K2B* alterations across TGCA pancancer datasets. **(C)** Heatmap of mevalonate pathway genes and PI5P4Ks in TGCA BRCA dataset. Subtypes are separated by PAM50 identification. *TP53* status (deletion=black bar, mutation=green bar, WT=gray bar). Heatmap is mRNA expression scaled by Z-score relative to diploid samples. **(D)** PIP4K2A expression in BRCA samples from TNMPlot using paired normal tissue compared to tumor. N=70 per group. Significance tested by paired t-test. p=**0.0084. **(E)** cBioPortal data of SQLE and DHCR7 mevalonate pathway gene expression, RSEM batch normalized from Illumina HiSeq_RNASeqV2. Clinical breast cancer subtypes are compared to normal samples from TCGA PanCancer BRCA dataset. Significance was tested using Kruskal-Wallis test with Dunn’s correction. p<0.01**, 0.001***.

**Figure S2.**
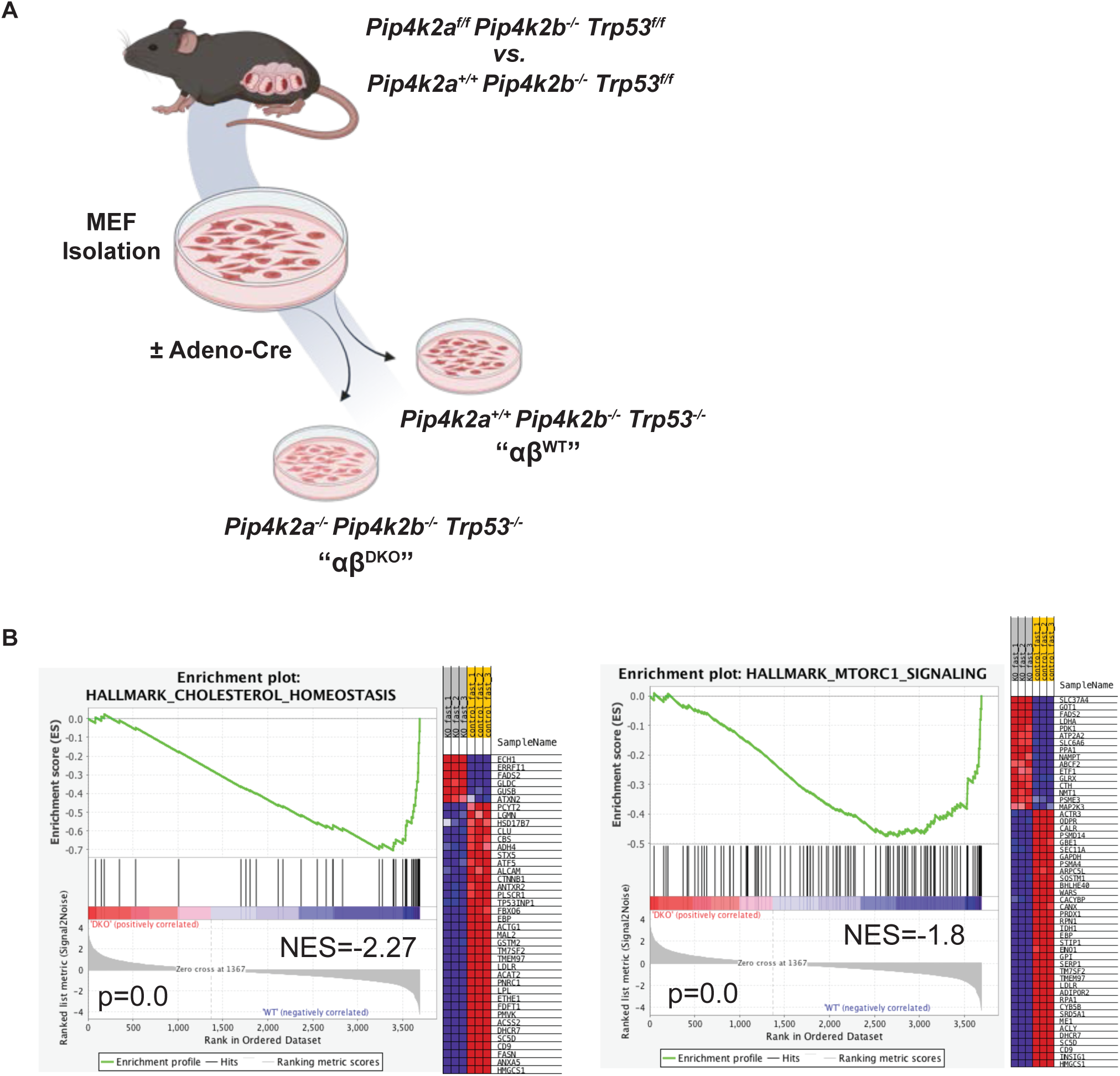
**(A)** Schematic of MEF generation from Trp53-/- mice. **(B)** RNA-sequencing of Pip4k2aflx/flx Pip4k2b-/- mice with adenoviral Cre (Pip4k2a-/- Pip4k2b-/- = DKO) or adenovirus empty vector (Pip4k2aflx/flx Pip4k2b-/- = WT). GSEA plots of cholesterol homeostasis and mTORC1 signaling from the Hallmarks MSigDB.

**Figure S3.**
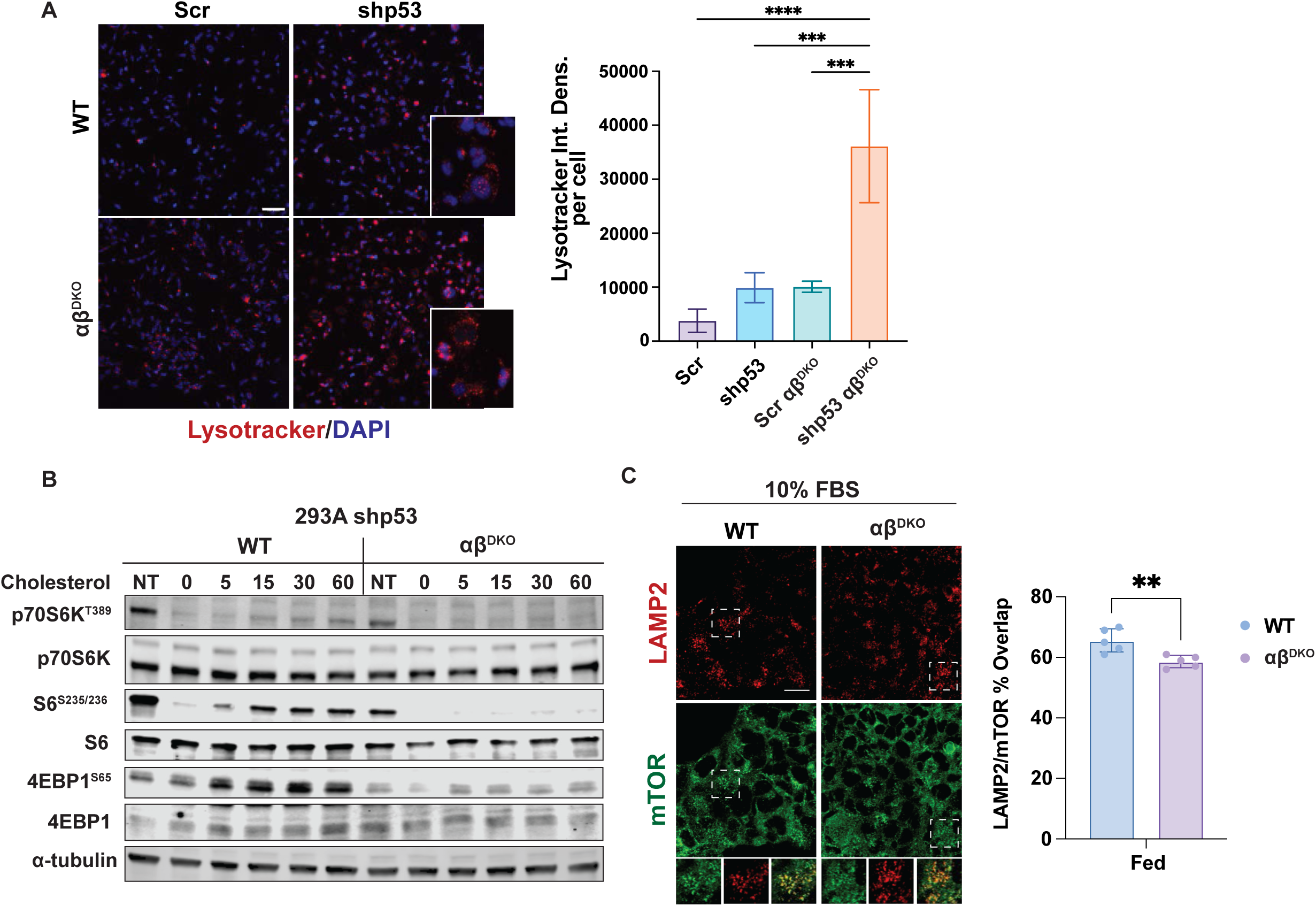
**(A)** Lysotracker staining of 293A shp53 cells with functional p53 (Scr), deficient p53 (shp53), functional p53 with PI5P4Kαβ DKO (αβDKO Scr) and deficient p53 with PI5P4Kab DKO (αβDKO shp53). Statistics represent two-way ANOVA with multiple comparisons compared with control (WT). Dunnett’s test used for multiple comparison correction, p-value <0.05*, 0.01**, 0.001***. **(B)** Immunoblot of 293A shp53 cells for PI5P4Ks and mTORC1 target genes in cholesterol depletion and repletion assay over a 60 minute time course. **(C)** 293A shp53 cells costained for endogenous mTOR and LAMP2. Scale bar=10uM Shown is the mean+SD of the calculated overlap. ANOVA with Bonferroni correction for multiple comparisons. p=****0.0012.

**Figure S4.**
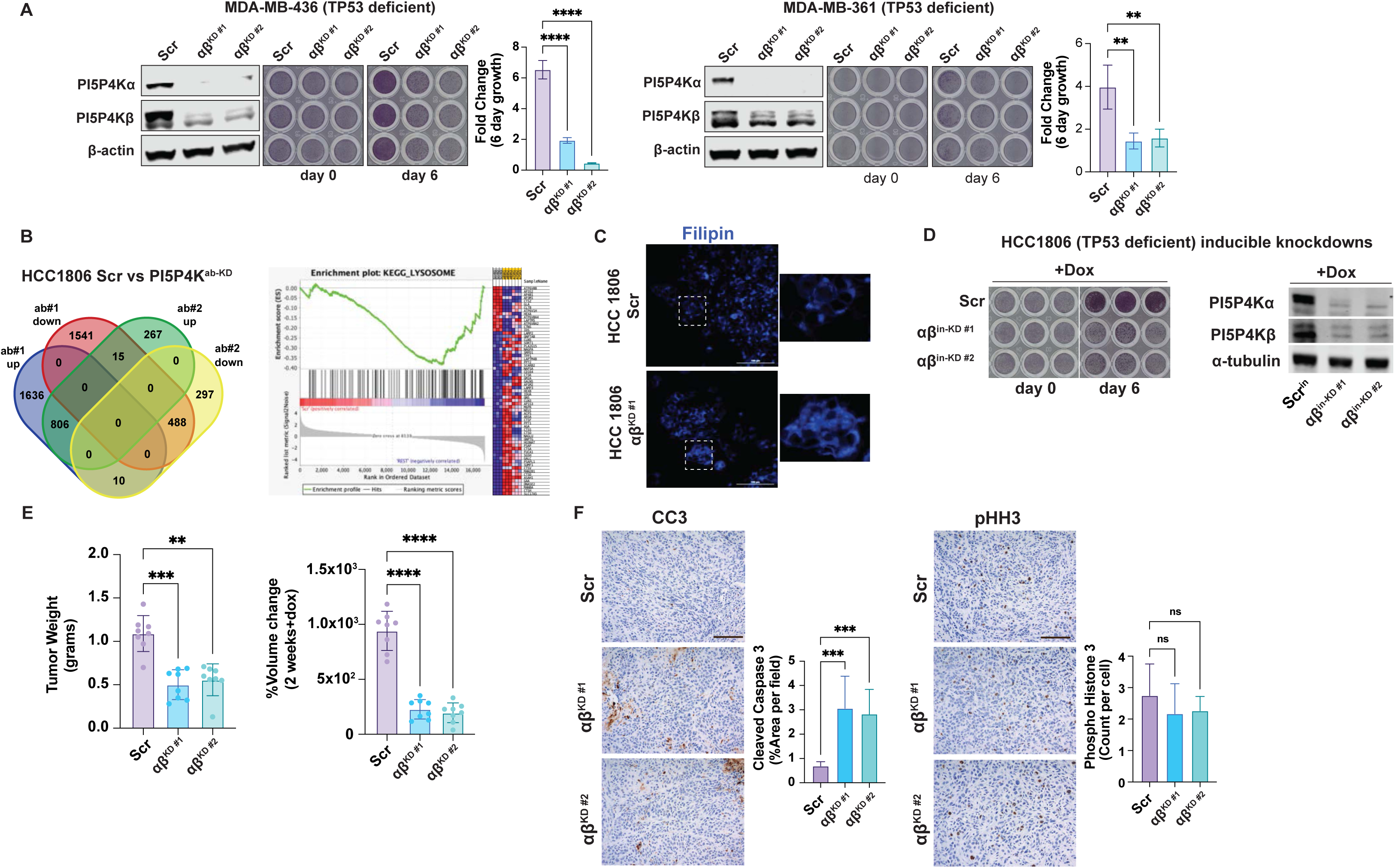
**(A)** Proliferation assays were performed using crystal violet on indicated cell lines with either non-targeting scramble shRNA (Scr) or dual, stable shRNA knockdown of PI5P4Ks (αβKD#1 or αβKD#2). N=3 independent replicates. Shown is mean + SD. ANOVA followed by Dunnett’s multiple comparison correction compared to Scr. p=*0.05, **0.01, ***0.001, ****0.0001. **(B)** *(left)* Venn diagram of overlapping significantly changed transcripts from HCC1806 RNA sequencing. Comparisons are scramble shRNA (Scr) vs either PI5P4Ks (αβKD#1) or PI5P4Ks (αβKD#2). *(right)* GSEA results from HCC1806 RNA sequencing of lysosomal genes. NES=(-)1.69, p=0.002. **(C)** Filipin staining of HCC1806 cells performed at 20x magnification. Scale bar = 100um. **(D)** Validation of inducible hairpins in HCC1806 cells. **(E)** HCC1806 tumor weights and percent volume change at experimental endpoint. Shown is a bar plot of mean+SD. ANOVA with multiple comparisons to Scr with Sidak correction. p=*0.05, **0.01, ***0.001, ****0.0001. **(F)** Immunohistochemistry of HCC1806 tumors stained for cleaved-caspase 3 (CC3) and phosphohistone-H3 (pHH3). n=10 fields per each subset. Significance was calculated by one-way ANOVA with Dunnett’s correction. p<*0.01, **0.001, ***0.0001. Scale bar = 500um

